# CD14 Blockade Attenuates Arrhythmogenic Cardiomyopathy

**DOI:** 10.64898/2026.09.30.755828

**Authors:** Vinay R. Penna, Emily A. Ehle, Waleed Farra, Elisa N. Cannon, Maicon Landim-Vieira, Ann M. Center, Macee C. Owen, Rajiu Venkatesan, Gyu Seong Heo, Avash Das, Yongjian Liu, Mark. W. Appleby, David. T. Crowe, Brian. W. Ziegelaar, Garry. L. Redlich, Jeffrey E. Saffitz, Stephen P. Chelko, Kory J. Lavine

## Abstract

**Background:** Arrhythmogenic cardiomyopathy (ACM) is a genetic and phenotypically heterogeneous myocardial disorder characterized by progressive cardiomyocyte loss, immune cell infiltration, and fibro-fatty replacement of the ventricular myocardium, with an increased risk of sudden cardiac death (SCD), predominantly caused by mutations in cardiac desmosomal genes including *PKP2, DSP*, and *DSG2.* We previously described the pivotal role of NFκB and CCR2^+^ cardiac macrophage-triggered inflammation in ACM pathogenesis.

**Methods:** WT or *Dsg2*^mut/mut^ mice were treated with either isotype control or anti-CD14 antibody once a week for the described treatment period (4 or 8 weeks). At the end of the treatment period, disease characteristics were assessed by echocardiography, electrocardiography, histology, immunofluorescence, positron emission tomography (PET) imaging and single-cell transcriptomics.

**Results:** Here, we demonstrate the efficacy of neutralizing CD14 signaling, an important mediator of inflammatory macrophage activation, in attenuating ACM disease *in vivo*. Additionally, we elucidate shifts in the cellular and transcriptomic landscapes that occur following anti-CD14 antibody treatment in a well-established mouse model of ACM and identify substantial reductions in inflammatory macrophage and pro-fibrotic fibroblast populations.

**Conclusion:** This study provides evidence that blockade of CD14 can attenuate the ACM disease phenotype. It may do so by inhibiting macrophage-based activation and inflammation, while also altering the transcriptional state of other important cell types. Collectively, these findings suggest that CD14 inhibition may be an effective option for ACM, which should be explored in early-phase clinical studies.

## Introduction

Arrhythmogenic cardiomyopathy (ACM) is a genetic form of nonischemic heart disease, affecting 1:2,000 to 1:5,000 people globally, although frequent misdiagnosis as myocarditis may underscore its prevalence.^1^ ACM is caused primarily by mutations in cardiac desmosomal genes such as plakophilin-2 (*PKP2*), desmoplakin (*DSP*), and desmoglein-2 (*DSG2*).^2,3^ It is one of the leading causes of sudden cardiac death due to fatal ventricular arrhythmias.^2,4^ Many individuals with ACM also progress to end-stage heart failure with heart transplantation being the only curative therapy. Other treatments, such as antiarrhythmics, standard medical therapies for heart failure, and implantable cardioverter defibrillators (ICDs) only temporize symptoms.^5,6^ Gene therapy, which involves replacement of defective genes with a healthy copy, has been proposed as a cure for ACM.^7^ While proof of concept experiments have been performed in preclinical models of ACM and a similar approach has been approved for other conditions such as Duchenne muscular dystrophy, there is still no approved gene therapy treatment for ACM.^8,9^

Extensive myocardial inflammation and fibrosis have been observed in over 70 percent of postmortem explants from ACM patients.^2^ Additionally, patients with ACM display elevated levels of inflammatory cytokines in their serum relative to healthy individuals.^2,4,10,11^ We previously demonstrated the importance of NFκB signaling in cardiac myocytes from two different iPSC-CM lines with pathogenic variants in *PKP2*c.^1851del123^ and D*SG2*^c.2358delA12^ in mediating ACM disease progression as well as the recruitment of C-C motif chemokine receptor 2 positive (CCR2^+^) monocytes and macrophages.^13,14^ Suppressing CCR2^+^ macrophage recruitment or neutralizing one of their most prominent cytokine products, interleukin-1β (IL1β), significantly attenuated ACM pathology, highlighting the potential benefits of anti-inflammatory therapies for this disease.^13–16^ These proof-of-principle studies set the stage for therapeutics that modulate immune activation and inflammation in ACM.

CD14 is an obligate co-receptor for lipopolysaccharide (LPS) and Toll-like receptor 4 (TLR4).^17^ In the setting of sterile immune activation, CD14 is required for sensing of damage-associated molecular patterns (DAMPs), released from dying cells, and is a major component of innate immune activation.^18^ Recently, CD14 has been used as a marker for autoimmune and inflammatory diseases.^19–21^ Furthermore, CD14 has been increasingly implicated in cardiovascular diseases, including atrial fibrillation; however, its involvement in ACM has been largely unexplored.^22–25^ Here, we demonstrate that CD14 is activated in ACM and that its inhibition decreased myocardial inflammation, diminished pathogenic fibrosis, favorably shifted cellular and transcriptomic landscapes of ACM, and attenuated ACM pathogenesis.

## Methods

A detailed methods section can be found in the Supplemental Appendix.

### Animals

All experiments conformed to the Guide for the Care and Use of Laboratory Animals from the National Institutes of Health (NIH publication no. 85–23, revised 1996). All animal protocols were approved by the Florida State University and Washington University in St. Louis Animal Care and Use Committees. Mice were housed in temperature- (20–22°C) and humidity-controlled rooms (40–60%), with a 12-hour light/dark cycle, and provided access to standard rodent chow and water *ad libitum*. Age-matched C57BL/6 mice served as WT controls. The generation of homozygous *Dsg2* mutant (*Dsg2*^mut/mut^) mice has been previously described.^14^

### CD14 antibody and treatment

A mouse anti-mouse antibody to CD14 (biG53 LALA-PG mAb) was provided by Implicit Bioscience Ltd. WT or *Dsg2*^mut/mut^ mice were treated (via intraperitoneal injections) with either anti-CD14 (αCD14) or isotype control (mouse IgG2a LALA-PG antibody, IgG) at a dose of 5 mg/kg/week for the following timelines: (a) 6 to 10 weeks, (b) 8–16 weeks, or (c) 16–24 weeks. biG53 LALA-PG mAb is a CD14 function-blocking antibody that does not deplete monocytes (unpublished data, Implicit Bioscience Ltd.). Previous PK/PD studies demonstrated that this dose amount, frequency, and route of administration of biG53 saturates monocyte CD14 for the entire dosing interval.

### Statistical Analysis

All data are expressed as mean±SEM (n-values inset within figure legends) for all *in vivo* studies. All data were normally distributed and were analyzed using parametric tests. Brown-Forsythe and Welch ANOVA were used for comparisons between multiple groups with unequal variances (unless noted otherwise within figure legends). P < 0.05 was deemed significant. All statistical analyses were done using GraphPad Prism (v10) software.

## Results

### CD14 expression and activation are increased in human ACM and *Dsg2*^mut/mut^ mice

CD14 is a TLR co-receptor for pathogen-associated molecular patterns (PAMPs) and DAMPs and is primarily expressed by myeloid cells, particularly monocytes and macrophages, where it drives inflammatory signaling.^18^ CD14, including its membrane associated and soluble forms, has become increasingly recognized as an elevated biomarker in cardiac diseases, including atrial fibrillation, and has been implicated in genome-wide association studies (GWAS) for cardiovascular risk.^22–24^ To determine if CD14 levels are increased in ACM, we performed flow cytometry using six-week-old WT and *Dsg2*^mut/mut^ mouse hearts, looking for increases in CD14^+^ myeloid populations (**Supplemental Figure 1**). Our prior work demonstrated *Dsg2*^mut/mut^ mice show increased arrhythmias and reduced cardiac function by 8 weeks of age,^13–15^ in the absence of observable ventricular fibrosis, thus six-week-old mice were selected to determine if an underlying inflammatory pathway was present prior to overt remodeling.

We observed an increase in CD14^+^ cells per milligram of heart tissue in *Dsg2*^mut/mut^ mice compared to WT controls (**Figure 1A, B**). To determine if there was an upregulation of active CD14 signaling in *Dsg2*^mut/mut^ mice, we used our previously generated mouse single-cell RNA sequencing data to assess differential expression of *Cd14* and the following genes associated with downstream CD14 signaling: *Nfkbiz, Tlr4, Myd88, Irak1, Irak4, Ikbkb, IL1b,* and *Ly6e.*^16,18^ We found the majority of these genes were increased in *Dsg2*^mut/mut^ mice relative to WT controls (**Supplemental Figure 2A**). Using these genes, we constructed a CD14 activation signature and found that it was predominantly expressed in myeloid cell types (**Supplemental Figure 2B**). Next, we turned to our previously generated human ACM single-nuclei RNA sequencing dataset.^16^ We found CD14 expression to be highest in myeloid cells, which also corresponded with elevated levels of NFκB transcription factor activity (predicted using DoRothEA) in human ACM myocardium (**Figure 1C, D**).^26^ We then converted our murine CD14 activation signature using corresponding human orthologs and found that it was increased in ACM cardiac explants relative to controls (**Supplemental Figure 2C**) and, as observed in *Dsg2*^mut/mut^ mice, elevated levels of NFκB transcription factor activity was additionally enriched in the myeloid population (**Supplemental Figure 2D**). These findings support the possibility that inhibiting CD14 signaling may reduce inflammation in ACM.

**Figure 1.**
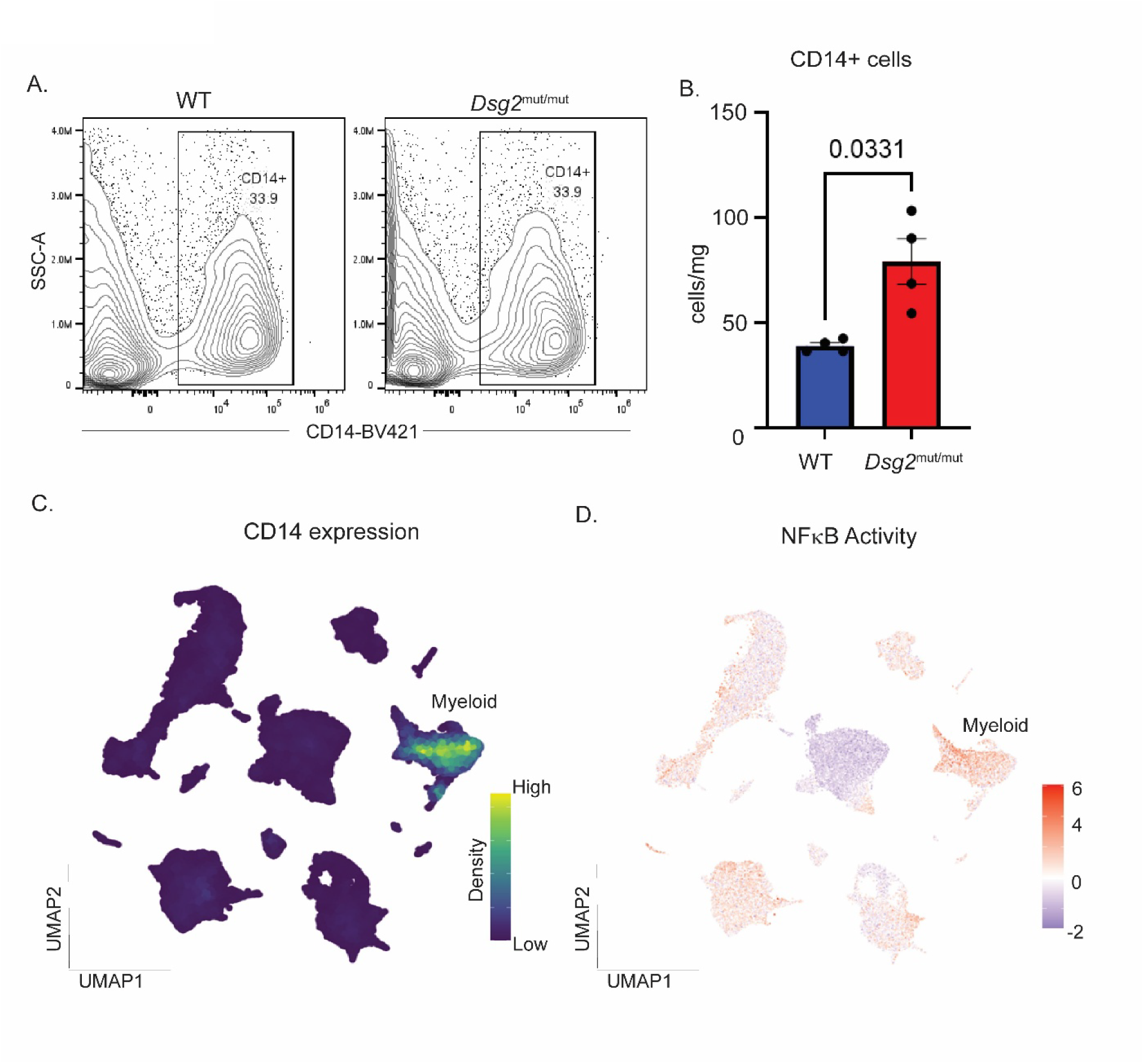
CD14 activity is elevated in both mouse and human ACM. (**A**) Representative flow cytometry plots of CD14^+^ cells in six-week-old WT and *Dsg2*^mut/mut^ mice. (**B**) Quantification of CD14^+^ cells per milligram of heart tissue. N=4 for each genotype. Significance determined by unpaired t-test with Welch correction. Error bars represent SE of the mean. (**C**) Density plot displaying *CD14* expression across our human ACM UMAP. (**D**) NFκB transcriptional activity displayed on our human ACM UMAP.

### Anti-CD14 antibody treatment decreases inflammatory macrophage infiltration in *Dsg2*^mut/mut^ mice

We treated six-week-old WT and *Dsg2*^mut/mut^ mice with either mouse αCD14 antibody or isotype control intraperitoneally once a week for four weeks until mice were ten weeks of age (**Figure 2A**). Our previous work indicated the contribution of CCR2^+^ macrophages in ACM pathogenesis, suggesting decreasing their numbers would reduce disease severity.^13,16^ Following treatment, we assessed the impact of CD14 inhibition on CCR2^+^ macrophage abundance in the heart. Using a CCR2-targeted PET radiotracer, we observed a significant decrease in CCR2 signal in the hearts of αCD14 treated *Dsg2*^mut/mut^ mice compared to those treated with isotype control (**Figure 2B**).

**Figure 2.**
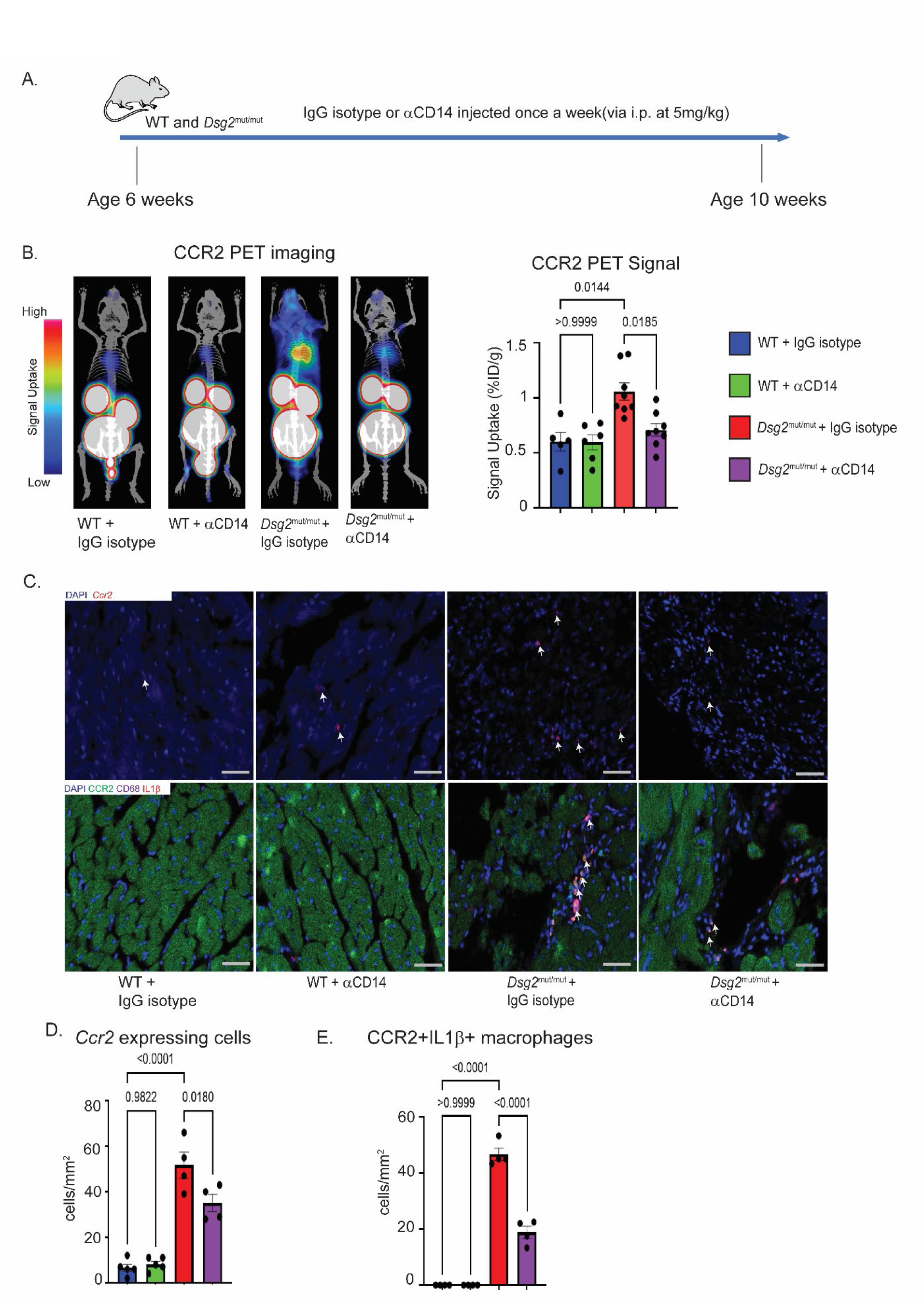
*α*CD14 treatment decreases inflammatory macrophage infiltration in *Dsg2*^mut/mut^ mice. (**A**) Experimental setup for αCD14 treatment. (**B**) Representative images of CCR2 PET imaging and quantification of CCR2 PET signal. N=5, 6, 8, and 8 for each group, respectively. Significance determined by Brown-Forsythe and Welch ANOVA with p-values inset. Error bars represent SE of the mean. (**C**) Representative *Ccr2 in situ* RNA hybridization (above) and CCR2, CD68, IL1β immunostained myocardium (below). Color coding is inset. White arrows indicate cells of interest: CCR2^+^ cells (above) and triple-labeled cells (CCR2^+^ CD68^+^ IL1β^+^) (below). (**D**) Quantification of CCR2^+^ cells from *in situ* hybridization. N=5, 5, 4, and 4 per group, respectively. Significance determined by one-way ANOVA with p-values inset. Error bars represent SE of the mean. (**E**) Quantification of CD68+CCR2^+^ IL1β^+^ cells from immunofluorescence data. N=5, 5, 4, and 4 per group, respectively. Significance determined by one-way ANOVA with p-values inset. Error bars represent SE of the mean.

Upon harvesting the hearts from these mice, we performed *in situ* RNA hybridization for *Ccr2* and found decreased numbers of *Ccr2*-expressing cells in *Dsg2*^mut/mut^ mice treated with αCD14 antibody (**Figure 2C, D**), consistent with our PET imaging data. Additionally, we assessed the presence of inflammatory macrophages following treatment and performed immunofluorescence staining for CD68, CCR2, and IL1β and found a substantial reduction in double-labeled CCR2^+^ IL1β^+^ cells in αCD14 treated *Dsg2*^mut/mut^ mice relative to isotype-treated counterparts (**Figure 2C, E**). Additionally, we performed multiplex cytokine array analyses and found that anti-CD14 antibody treatment decreased myocardial levels of several pro-inflammatory and pro-fibrotic cytokines, including IL1β, CXCL2 (MIP2α), CXCL9 (MIG), osteopontin (OPN), and periostin (POSTN) (**Table 1**).

**Table 1.** Major pro-inflammatory and pro-fibrotic cytokines.

| Cytokine | WT + αIgG vs *Dsg2*<sup>mut/mut</sup> + αIgG | | WT + αIgG vs *Dsg2*<sup>mut/mut</sup> + αCD14 | |
| --- | --- | --- | --- | --- |
|  | Fold Change | P-value | Fold Change | P-value |
| CCL17 | 3-5 | 0.087 | 1-3 | 0.894 |
| CD14 | 5-8 | $\leq 0.01$ | 3-5 | 0.149 |
| CD40 | 3-5 | $\leq 0.05$ | $\leq 1$ | 0.944 |
| CHI3L1 | $\geq 12$ | $\leq 0.01$ | 1-3 | 0.764 |
| C5/C5a | 8-12 | $\leq 0.01$ | 1-3 | 0.833 |
| CRP | 3-5 | $\leq 0.01$ | 1-3 | 0.287 |
| CXCL9 (MIG) | 3-5 | $\leq 0.01$ | $\leq 1$ | 0.971 |
| CXCL2 (MIP2α) | 5-8 | $\leq 0.01$ | 1-3 | 0.622 |
| CXCL13 | 8-12 | 0.071 | $\leq 1$ | 0.833 |
| GDF15 | 3-5 | 0.099 | $\leq 1$ | 0.927 |
| IL-1α | 3-5 | 0.109 | $\leq 1$ | 0.849 |
| IL-1β | $\geq 12$ | $\leq 0.01$ | 1-3 | 0.150 |
| IL-17a | $\geq 12$ | $\leq 0.01$ | $\leq 1$ | 0.274 |
| OPN | $\geq 12$ | $\leq 0.01$ | 8-12 | $\leq 0.01$ |
| POSTN | $\geq 12$ | $\leq 0.01$ | 5-8 | 0.199 |

### Anti-CD14 antibody treatment alters the transcriptional profile of myeloid populations in *Dsg2*^mut/mut^ mice

To further elucidate the impact of αCD14 antibody treatment on the overall myeloid population, we performed single-cell RNA sequencing on ten-week-old WT and *Dsg2*^mut/mut^ mice, following four weeks of treatment with either αCD14 antibody or isotype control as described above. We sorted both immune and fibroblast populations from these treated mouse hearts at 10 weeks of age to generate separate libraries. Following sample quality control and normalization, we divided the immune library into myeloid and lymphoid populations (**Supplemental Figure 3A**). Sub-clustering analysis revealed 14 distinct myeloid populations: (1) dendritic cells (DC): DC1 (*Xcr1, Clec9a*), DC2-1 (*Cd209a, Clec10a*), DC2-2 (*Klrd1, Bc11a*), migratory (Mig) MigDCs (*Ccr7, Ccl5*), and plasmacytoid (p) pDCs (*Siglech, Runx2*); (2) monocytes (Mono): Mono1 (*Plac8, Ly6c2*), Mono2 (*Thbs1, Ccl2*), and Mono3 (*Ace, Itgal*); and (3) macrophages (Mac): Mac1 (*Cxcl2, Cd14*), Mac2 (*C1qa, C1qc*), Mac3 (*Isg15, Ifit3*), Mac4 (*Top2a, Mki67*), Mac5 (*Cd207, Vsig4*), and Mac6 (*Retnla, Lyz1*) (**Figure 3A** and **Supplemental Figure 3B**). To determine which populations are relevant in cardiac diseases, we used our previously developed human ACM disease signature and overlaid it onto the mouse UMAP space (**Figure 3B**).^16^ This signature overlaid primarily with the following populations: Mac1, Mono1, and Mono2. The collective proportion of these populations was reduced in αCD14 treated *Dsg2*^mut/mut^ mice relative to isotype-treated controls (**Figure 3C**).

**Figure 3.**
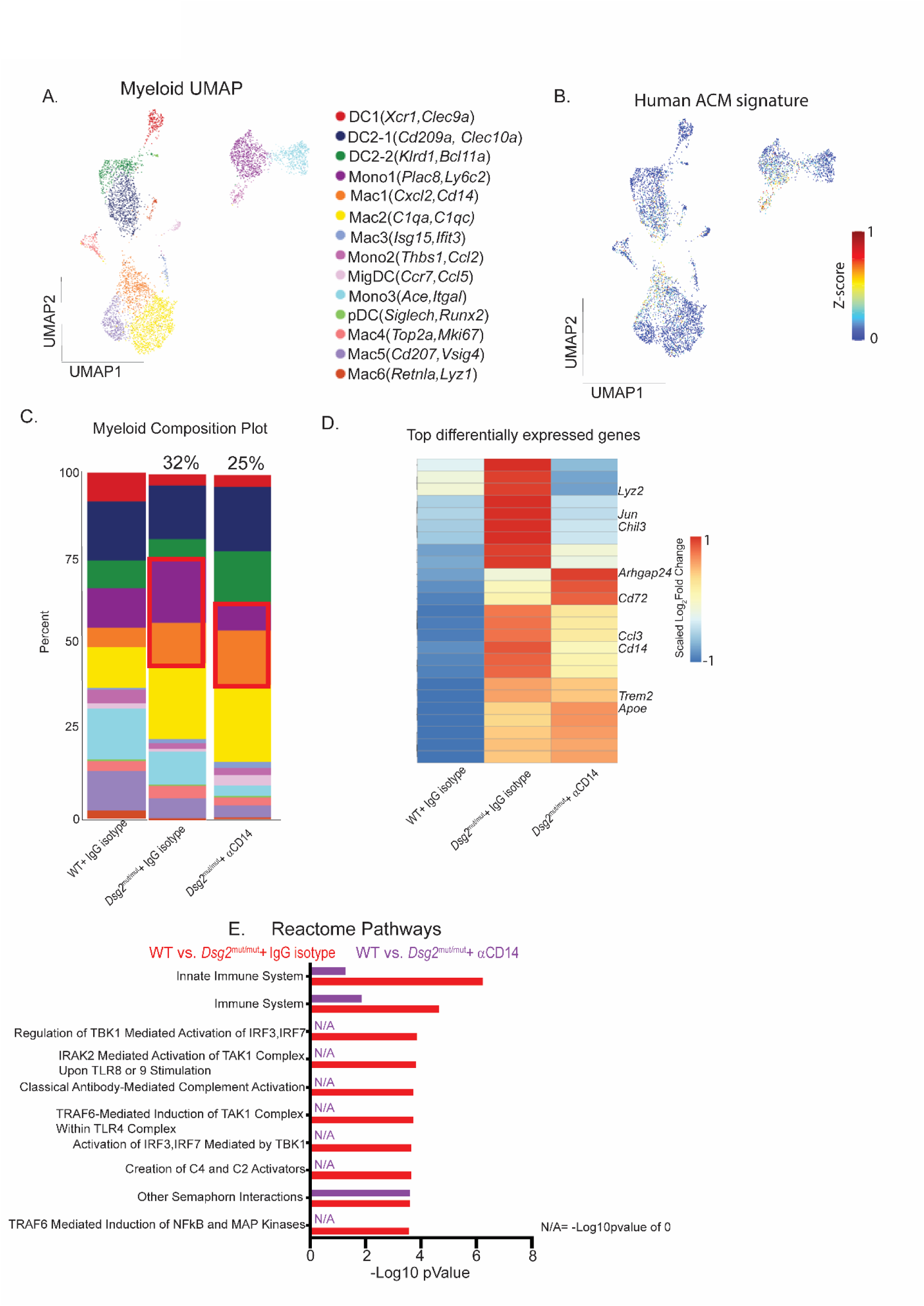
*α*CD14 antibody treatment alters the transcriptional profile of myeloid populations. (**A**) Global UMAP of myeloid populations. (**B**) Gene expression score generated from mouse orthologs of differentially upregulated myeloid genes from human ACM sequencing data overlaid onto mouse myeloid UMAP. (**C**) Composition plots comparing myeloid populations between isotype-treated WT and *Dsg2*^mut/mut^ mice, and αCD14 treated *Dsg2*^mut/mut^ mice. (**D**) Heatmap displaying top differentially expressed myeloid genes between isotype-treated WT and *Dsg2*^mut/mut^ mice compared to those same genes in αCD14 treated *Dsg2*^mut/mut^ mice. (**E**) Reactome biological pathways for the top 25 differentially upregulated genes in WT vs *Dsg2*^mut/mut^ isotype treated mice (red) and WT vs *Dsg2*^mut/mut^ αCD14 treated mice (purple).

Next, we determined if the broader ACM phenotype had been reversed in myeloid cells following αCD14 antibody treatment. To address this question, we determined the expression values of the top 25 differentially expressed genes between WT and *Dsg2*^mut/mut^ isotype control mouse hearts and assessed whether these genes were impacted by αCD14 antibody treatment in *Dsg2*^mut/mut^ hearts (**Figure 3D**). We observed that a variety of genes upregulated in ACM myeloid cells were downregulated following αCD14 antibody treatment, including genes associated with inflammation and immune activation (*Jun, Chil3, Lyz2, Ccl3*).^27–29^ Among genes that were upregulated following αCD14 treatment, we observed increased expression of *Arhgap24*, a negative regulator of NFκB signaling, *Apoe*, known to shift pro-inflammatory macrophages to an anti-inflammatory macrophage state,^30^ and *Cd72*, a negative regulator of immune cell activation and IFNγ production,^31^ further suggesting the anti-inflammatory effects of targeting CD14.^32^ Next, we performed pathway enrichment analysis using differentially expressed genes between WT and *Dsg2*^mut/mut^ isotype controls. We observed an increase in pathways associated with innate immune activation and NFκB signaling (**Figure 3E**). Enrichment of these pathways was additionally reduced when comparing WT mice to αCD14 treated *Dsg2*^mut/mut^ mice (**Figure 3E**). These findings suggest that αCD14 treatment reduces inflammatory macrophage populations and transcriptional activity associated with innate immune activation.

Upon further examining the myeloid clusters, we noticed a difference in DC populations. To further explore these findings, we sub-clustered the DCs and identified 7 distinct populations: classical DC1 (*Xcr1, Irf8*); two groups of classical DC2s: DC2-1 (*Atf3, Ier3*) and DC2-2 (*Arhgap24, Sirpb1a*); Interferon gene (IFN) expressing DCs (*Cxcl9, Mt1*); migratory DCs (MigDC) (*Ccr7, Mreg*); monocyte DCs (moDC) (*Lyz2, S100a6*); and plasmacytoid DCs (pDC) (*Siglech, Runx2)* (**Supplemental Figure 4** and **Supplemental Figure 5A**). DC2-1 and IFN DCs were increased in *Dsg2*^mut/mut^ isotype controls relative to αCD14 treated *Dsg2*^mut/mut^ mice. Conversely, DC2-2 populations were expanded following αCD14 treatment (**Supplemental Figure 5B**). DCs are very heterogeneous cell types and have not been thoroughly characterized in the heart.^33^ As a result, we turned to the cancer literature to help identify important gene signatures in our populations.^34–37^ We constructed a pro-inflammatory/fibrotic signature consisting of *Cd40, Cd80, Cd86, Cd14, C1qa, Cd163, Cd68, Il18, Dennd4a*, and *Tnip3* genes, and a pro-resolving and immunoregulatory signature consisting of *Fgr, St8sia6, Ddr1,* and *Siglece* genes. The former signature was more robustly expressed in DC2-1 and IFN DC, the populations increased in *Dsg2*^mut/mut^ mice given isotype control, while the latter signature was increased in DC2-2, the population increased in *Dsg2*^mut/mut^ mice treated with αCD14 antibody (**Supplemental Figure 5C** and **Supplemental Figure 5D**). These findings suggest that αCD14 treatment can alter DC state to a more pro-resolving, immunoregulatory phenotype.

### Anti-CD14 antibody treatment restores the proportion of natural killer cells in the heart

We next proceeded to examine cardiac lymphoid populations. Clustering analysis revealed 11 distinct populations: activated CD4 T cells, activated CD8 T cells, B cells, gamma-delta T cells, mast cells, naïve CD4 T cells, naïve CD8 T cells, natural killer (NK) cells, plasma cells, stem-like CD8 T cells, and regulatory T cells (**Supplemental Figure 6** and **Supplemental Figure 7A**). The most prominent difference we observed was a decrease in the proportion of NK cells in isotype-treated *Dsg2*^mut/mut^ mice relative to WT controls and a restoration of that proportion in *Dsg2*^mut/mut^ mice treated with αCD14 antibody (**Supplemental Figure 7B**). We corroborated this finding via flow cytometry (**Supplemental Figure 7C** and **Supplemental Figure 8**). NK cells are believed to play a homeostatic and protective role in the heart through a variety of mechanisms,^38^ and have additionally been found to be reduced in peripheral blood samples from patients with ACM compared to healthy controls.^39^ Restoration of NK cell levels may be a sign of disease attenuation; conversely, their reduction may be an indicator of active disease onset and/or progression.

### Anti-CD14 antibody treatment alters the transcriptional state of fibroblasts in *Dsg2*^mut/mut^ mice

Given the importance of pathogenic fibrosis in heart failure and ACM disease progression, we explored transcriptional changes in fibroblast populations following αCD14 treatment.^40^ Following quality control, normalization, and clustering, we identified 9 fibroblast (Fib) populations: Fib1 (*Morrbid, Pla1a*), Fib2 (*Postn, Comp*), Fib3 (*Sntg1, Fgl2*), Fib4 (*Igfbp3, Cytl1*), Fib5 (*Opcml, Igfbp5*), Fib6 (*Cxcl14, Penk*), Fib7 (*Mmp3, Ccl19*), Fib8 (*Ptx3, Ccl2*), and Fib9 (*Ifit3, Isg15*) (**Supplemental Figure 9A** and **Supplemental Figure 9C**). Fib2 contained *Postn-*expressing fibroblasts, which we previously identified as a key disease-associated fibroblast population in ACM hearts.^13,16^ The proportion of Fib2 was increased in *Dsg2*^mut/mut^ mice relative to WT controls (**Supplemental Figure 9B**). To ascertain if there were any differences within this disease-associated fibroblast cluster between treatment groups, we divided this population into more deeply defined subsets. We identified 6 disease (dFib)-associated fibroblast clusters: dFib1 (*Ly6a, Tmem100*), dFib2 (*C4b, Sfrp2*), dFib3 (*Tagln, Acta2*), dFib4 (*Thbs4, Cilp2*), dFib5 (*Fmo2, Rbp4*), and dFib6 (*Ltbp1, Rgs5*) (**Supplemental Figure 10** and **Figure 4A**). We observed a reduction in dFib4 and a substantial increase in dFib3 in *Dsg2*^mut/mut^ mice treated with αCD14 antibody (**Figure 4B**). We noticed that the dFib4 population selectively expressed a number of genes previously associated with pathogenic, pro-fibrotic remodeling fibroblasts, including *Thbs4, Fap, Thbs1, Meox1, Runx1, Cilp2, Comp, Col1a1, Thy1, Isg15, Col3a1, Postn, Aebp1*, while dFib3 expressed genes associated with myofibroblasts, such as *Tagln, Acta2,* and *Svep1.*^41^ When comparing these gene signatures to our previously generated human ACM fibroblast disease signature, we found that the pro-fibrotic signature overlapped robustly with the human ACM disease signature. Conversely, this overlap was not observed with the myofibroblast signature (**Figure 4C**).^16^ These findings suggest that αCD14 treatment alters the transcriptional landscape of fibroblasts and shifts their phenotype towards a myofibroblast phenotype and away from a pathogenic lineage, specifically, fibroblast activation protein-positive (FAP^+^) fibroblasts. This would be consistent with our previous work examining the effects of IL1β neutralization on fibroblast cell state transitions.^41^

**Figure 4.**
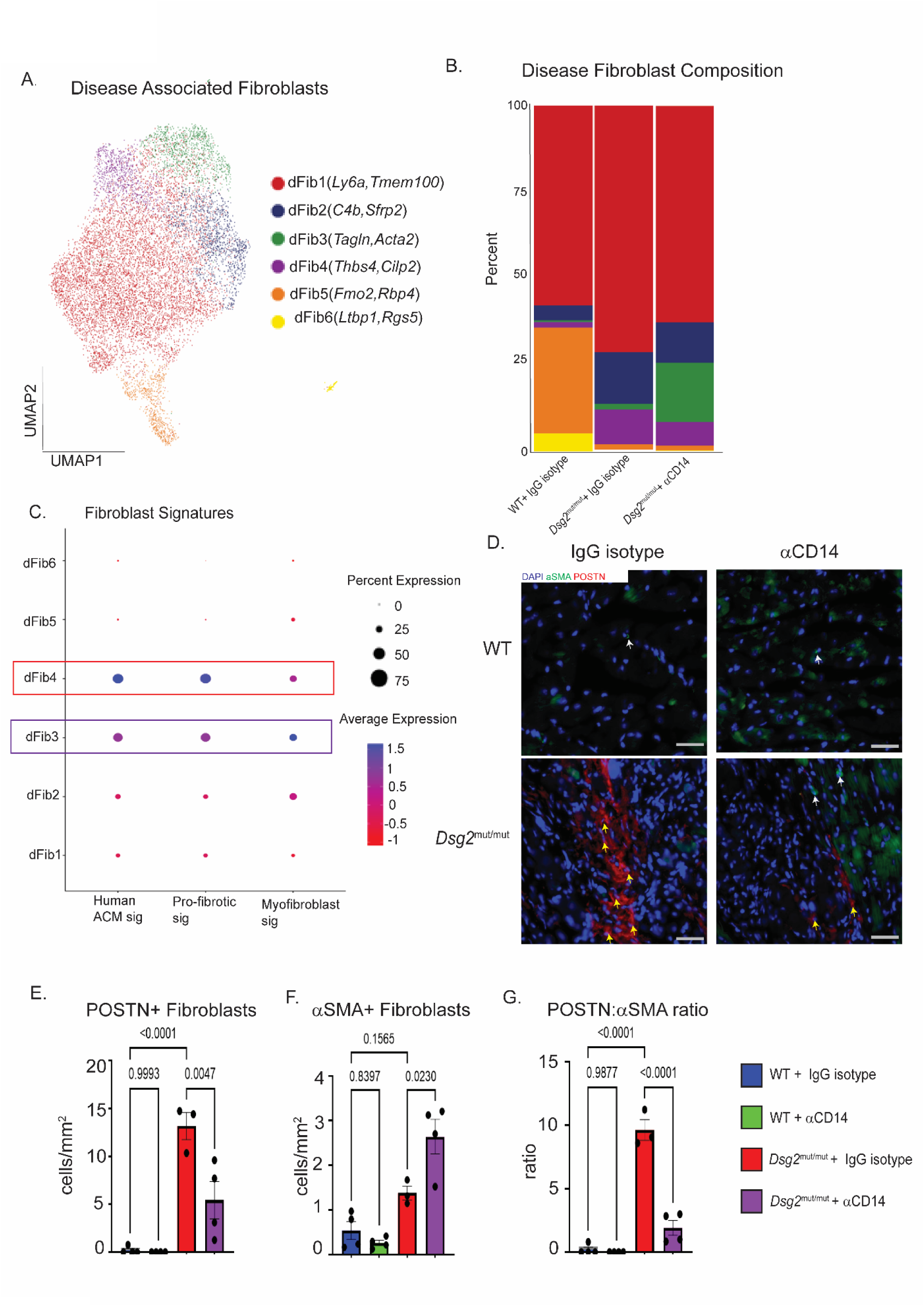
*α*CD14 antibody treatment alters the transcriptional state of fibroblasts. (**A**) UMAP of disease-associated fibroblasts. (**B**) Composition plots comparing disease-associated fibroblast populations between isotype-treated WT and *Dsg2*^mut/mut^ mice, and αCD14 treated *Dsg2*^mut/mut^ mice. (**C**) Dot plot comparing the human ACM fibroblast signature with the pro-fibrotic and myofibroblast signatures across all disease-associated fibroblast populations. (**D**) Representative images of POSTN and 〈SMA immunostained myocardium. Color code is inset. White arrows mark 〈SMA^+^ cells and yellow arrows mark POSTN^+^ cells. (**E**) Quantification of POSTN^+^ cells. N=5, 4, 3, and 4 per group, respectively. Significance determined by one-way ANOVA with p-values inset. Error bars represent SE of the mean. (**F**) Quantification of 〈SMA^+^ cells. N=5, 4, 3, and 4 per group, respectively. Significance determined by one-way ANOVA with p-values inset. Error bars represent SE of the mean. (**G**) Quantification of POSTN^+^ to 〈SMA^+^ cell ratio. N=5, 4, 3, and 4 per group, respectively. Significance determined by one-way ANOVA with p-values inset. Error bars represent SE of the mean.

We further corroborated our single-cell RNA sequencing findings by performing immunofluorescence staining for POSTN^+^ and α-smooth muscle actin-positive (αSMA^+^) levels in fibroblasts, which are representative markers for the pro-fibrotic fibroblast population and the myofibroblast population, respectively (**Figure 4D**). We observed that POSTN^+^ fibroblasts were indeed decreased in αCD14 treated *Dsg2*^mut/mut^ hearts, while αSMA^+^ fibroblasts were increased, and the overall POSTN to αSMA ratio was substantially decreased with αCD14 antibody treatment (**Figure 4E-G**).

### Anti-CD14 treatment attenuates disease characteristics in *Dsg2*^mut/mut^ mice

We modified our αCD14 treatment experiment, beginning treatment at eight weeks of age and continuing until sixteen weeks of age, with the intent to assess major disease characteristics in *Dsg2*^mut/mut^ mice (**Figure 5A**). At 8 weeks of age, *Dsg2*^mut/mut^ mice show mild dysfunction and arrhythmias; whereas by 16 weeks of age they show cardiac dysfunction and malignant ventricular arrhythmias, with overt cardiac remodeling and biventricular myocardial inflammation and fibrosis.^13–15^ Prior to and following antibody treatment, we performed echocardiography. We observed significantly higher left ventricular ejection fraction in αCD14 treated *Dsg2*^mut/mut^ mice relative to *Dsg2*^mut/mut^ mice that received isotype control, demonstrating a preservation of ejection fraction during treatment (**Figure 5B-C**). We also observed a decrease in the frequency of ectopic beats in αCD14 antibody treated *Dsg2*^mut/mut^ mice compared to isotype controls (**Figure 5D**). Following treatment, hearts were harvested and assessed for myocardial fibrosis, in which a significant decrease in myocardial fibrosis was noted in αCD14 antibody treated *Dsg2*^mut/mut^ mice compared to isotype-treated counterparts (**Figure 5E** and **Figure 5F**).

**Figure 5.**
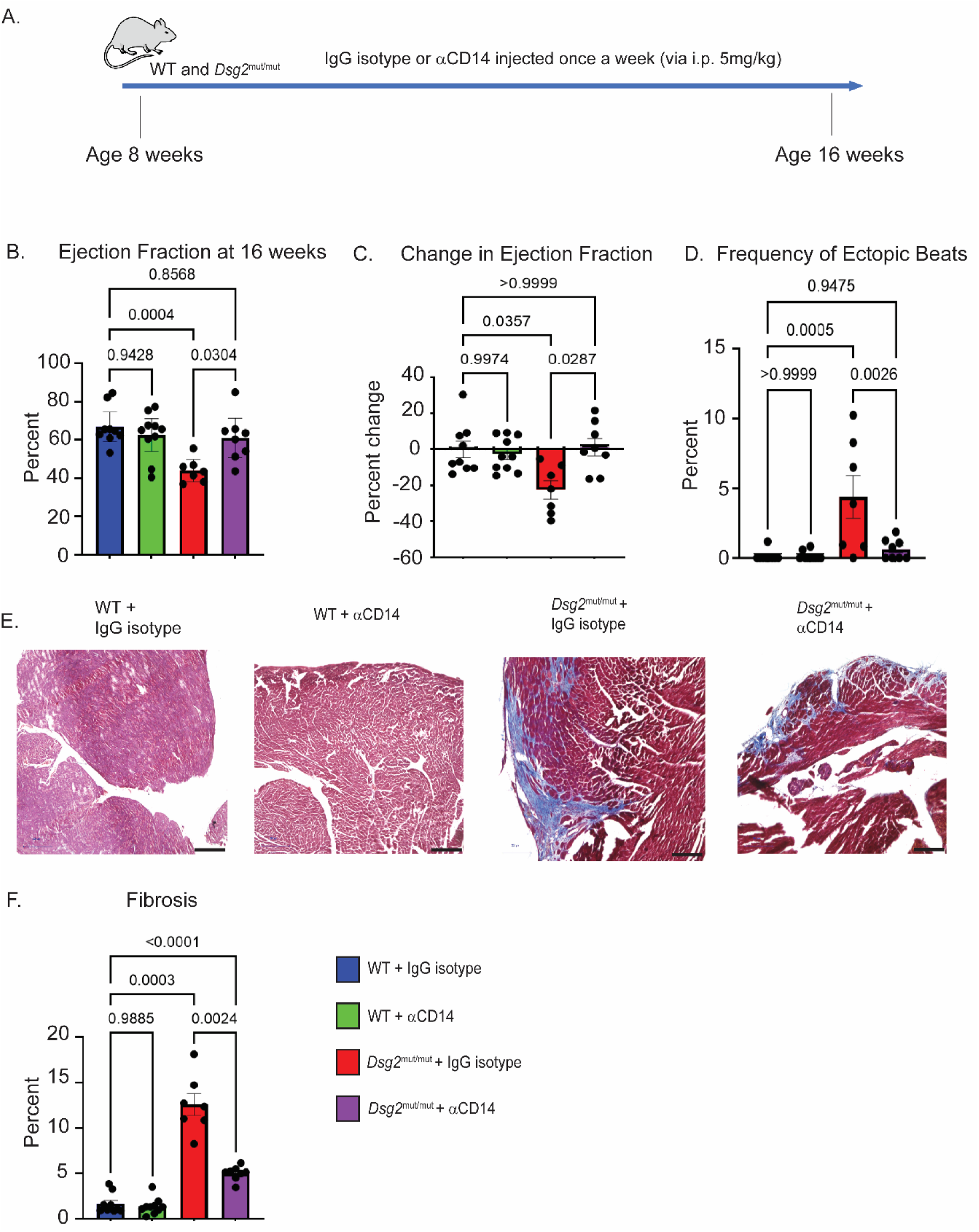
*α*CD14 treatment attenuates disease characteristics in *Dsg2*^mut/mut^ mice. (**A**) Experimental setup for αCD14 treatment. (**B**) Quantification of percent left ventricular ejection fraction (%LVEF) at 16 weeks. N=9, 10, 7, and 8 for each group, respectively. Significance determined by Brown-Forsythe and Welch ANOVA with p-values inset. Error bars represent SE of the mean. (**C**) Quantification of change of %LVEF at the start and end of treatment. N=9, 10, 7, and 8 for each group, respectively. Significance determined by one-way ordinary ANOVA with p-values inset. Error bars represent SE of the mean. (**D**) Quantification of percent ectopic beats (%PVCs) at 16 weeks. N=9, 10, 7, and 8 for each group, respectively. Significance determined by one-way ordinary ANOVA with p-values inset. Error bars represent SE of the mean. (**E**) Representative images of Masson’s trichrome stained hearts at 16 weeks. (**F**) Quantification of percent myocardial fibrosis at 16 weeks. N=9, 10, 7, and 8 for each group, respectively. Significance determined by Brown-Forsythe and Welch ANOVA with p-values inset. Error bars represent SE of the mean.

To assess whether a similar benefit could be derived from αCD14 antibody treatment at a later time point in which cardiac dysfunction and biventricular fibrosis are quite evident,^13–15^ we treated WT or *Dsg2*^mut/mut^ mice from 16- to 24 weeks of age (**Supplemental Figure 11A**). We observed that intervening at this stage of disease progression produced similar improvements in left ventricular ejection fraction (**Supplemental Figure 11B** and **Supplemental Figure 11C**), but not in the frequency of ectopic beats (**Supplemental Figure 11D**). Assessment of myocardial fibrosis showed substantial reduction in fibrosis following αCD14 antibody treatment, but to a much lesser extent compared to our prevention study (i.e., 8 to 16 weeks of age) (**Supplemental Figure 11E** and **Supplemental Figure 11F**). These results are indicative of the potential clinical benefits of αCD14 antibody treatment across the time course of ACM disease pathogenesis with the greatest benefit observed with early intervention.

## Discussion

Inflammation has long been recognized as a pathological hallmark of ACM; however, the role of inflammation and/or the various inflammatory mediators that contribute to ACM disease onset and progression have not been well characterized.^2,4,10,11^ Recently, we and others have demonstrated the active contribution of inflammation in ACM pathogenesis.^11,13,15^ Our work has shown the critical role of NFκB signaling in cardiomyocytes and CCR2^+^ monocyte-derived cardiac macrophages, as well as the primordial pro-inflammatory cytokine, IL1β, in driving ACM disease.^13,15,16^ These findings warrant the exploration of additional anti-inflammatory-based therapeutic approaches to expand the limited repertoire of treatments for ACM.

CD14, a key regulator in innate immune signaling and activation of inflammation, has been increasingly used as a prognostic marker for a variety of diseases.^20–23,25^ Additionally, polymorphisms in the gene have been associated with cardiovascular disease.^24^ Here, we observe that this finding extends to ACM as CD14^+^ cells are increased in the hearts of the *Dsg2*^mut/mut^ mouse model of ACM and that expression of genes associated with CD14 activation and downstream signaling are increased in both ACM mice and myocardial samples from patients with ACM.

To demonstrate the potential efficacy of CD14 blockade, we utilized a potent αCD14 antibody. Following treatment, we observed a drastic decline in CCR2^+^ macrophages via non-invasive whole heart PET imaging present in the hearts of Dsg2mut/mut mice, which we have previously shown is associated with disease attenuation.^13^ Additionally, we saw a decrease in CCR2^+^ macrophages actively producing IL1β, suggesting that blocking CD14 may reduce CCR2^+^ macrophage activation and inflammatory capabilities.

To gain further understanding of the transcriptional changes that may occur following αCD14 treatment in key immune and fibroblast populations, we performed single-cell RNA sequencing. While assessing the myeloid populations, we observed a reduction in the proportion of clusters associated with our signature of ACM disease in *Dsg2*^mut/mut^ mice treated with αCD14 relative to isotype-treated controls. Furthermore, expression of many genes, including those affiliated with inflammation, that were upregulated in isotype-treated *Dsg2*^mut/mut^ mice relative to WT controls were reduced in αCD14 treated *Dsg2*^mut/mut^ mice. Accordingly, there was a reduction in the enrichment of pathways associated with NFκB and other innate immune signaling cascades in *Dsg2*^mut/mut^ mice treated with αCD14.

Moving beyond macrophages, we observed a transcriptional shift in dendritic cells towards a more pro-resolving and immunoregulatory phenotype^34–37^ and a restoration of cardiac natural killer cell levels. More intriguingly, we noticed a shift in fibroblasts away from a pathogenic pro-fibrotic population to a myofibroblast-like group. Our previous work demonstrated a similar shift following IL1β inhibition in a model of angiotensin-induced hypertension.^41^ In that work, this shift was associated with reduced myocardial fibrosis, a finding we additionally observed in this study in ACM.

Although we believe the changes observed in myeloid cell types were likely due to the direct action of the αCD14 antibody, the mechanism(s) behind the transcriptional shift in the other cell types is less clear. It is possible that this is mediated via a direct interaction between pro-inflammatory macrophages and other immune and stromal cells, as has been demonstrated in other disease settings.^41–44^ Another possibility is an indirect action mediated by an alteration of the cardiac microenvironment.^45^ Further work will be necessary to determine the exact mechanisms at play.

Finally, we observed robust phenotypic improvements and attenuation of ACM disease characteristics following αCD14 treatment. *Dsg2*^mut/mut^ mice treated with αCD14 antibody displayed greater left ventricular ejection fraction, lower frequency of ectopic beats, and reduced myocardial fibrosis. Although these findings were most evident following treatment initiation at an early stage of disease development (i.e., 8–16 weeks of age), there were still apparent improvements even at a later stage of disease (i.e., 16–24 weeks) following intervention with αCD14 antibody, suggesting clinical benefits can be achieved using this treatment over the course of the disease’s natural history.

### Study Limitations

While our findings demonstrate the benefits of αCD14 therapy in a mouse model of ACM, we recognize that not all aspects of human ACM disease are recapitulated in mouse models of inherited diseases. Additionally, our treatment strategy does not distinguish between membrane-bound and soluble forms of CD14. As a result, we were unable to determine whether there was a differential role between these two different forms of CD14 in ACM progression.

### Novelty and Significance

#### What is known

- Arrhythmogenic cardiomyopathy (ACM) is a genetic form of heart disease predominantly caused by mutations in cardiac desmosomal genes including *PKP2, DSP*, and *DSG2*.
- Arrhythmogenic cardiomyopathy (ACM) has a strong component of myocardial inflammation leading to extensive cardiomyocyte cell death and replacement fibrosis, serving as a substrate for ventricular arrhythmias and contributing to cardiac dysfunction.

### New information

- CD14 expression and activation are increased in both human ACM as well as in an established mouse model of the disease (*Dsg2*^mut/mut^ mice).
- Anti-CD14 antibody treatment reshapes the cellular and transcriptomic landscape of the myocardium with prominent effects on macrophages and fibroblasts, resulting in shifts away from pathogenic cell states.
- Inhibition of CD14 signaling using a neutralizing antibody leads to significant attenuation of ACM pathogenesis in the *Dsg2*^mut/mut^ mouse model of ACM.

Myocardial inflammation has been observed in ACM for some time. Recent work has established that this inflammation, particularly NFκB mediated innate immune signaling, contributes to the pathogenesis of ACM, opening new opportunities for disease identification, management and therapeutic strategies. In this current study, we further identify CD14 as a novel therapeutic target. We demonstrate that CD14 expression and downstream activation are increased in both myocardium from the Dsg2-mutant mouse model of ACM and in hearts from patients with ACM. Importantly, pharmacologic blockade of CD14 with an anti-CD14 antibody substantially attenuated disease progression, reducing CCR2+ inflammatory macrophage infiltration, production of Interlukin-1 β (IL-1β), and several other pro-inflammatory cytokines, chemokines, and fibroadipokines, and importantly, attenuated myocardial fibrosis, ventricular ectopy, and cardiac dysfunction. Our single-cell RNA sequencing data provided additional mechanistic insights, revealing that CD14 inhibition remodels the cardiac immune and stromal microenvironment. Treatment reduced inflammatory macrophage populations and NF-κB-associated transcriptional programs, shifted dendritic cells toward a more pro-resolving and immunoregulatory state, and restored cardiac natural killer cell abundance. Notably, CD14 blockade also altered fibroblast states, reducing pathogenic POSTN+/FAP-associated pro-fibrotic fibroblasts while increasing a myofibroblast-like population. Collectively, these findings establish CD14 as an upstream regulator of inflammatory and fibrotic remodeling in ACM and provide preclinical evidence that CD14 inhibition can modify disease progression, supporting its potential translational utility into early-phase clinical studies.

## Acknowledgements

We would like to thank Hannah Luehmann and Nikki Fettig for their assistance with the PET imaging studies. We thank the Genome Technology Access Center at the McDonnell Genome Institute at Washington University School of Medicine for their help with sequencing and genomic analysis. We thank Implicit Bioscience for generously providing the αCD14 antibody and isotype control antibody.

## Funding

American Heart Association Career Development Award 19CDA34760185 (To SPC); an Institute of Peadatric Rare Diseases (IPRDS) Award from FSU COM IPRDS (TO SPC), a Florida Heart Research Foundation (FHRF) Cardiovascular Doctoral Student Grant (to GHDNA), American Heart Association Predoctoral Fellowship Award 25PRE1372729 (to E.A.S.), an FSU Research Foundation Award (to SPC), a National Institutes of Health grant1F31HL181979-01 (to EAS), a National Institutes of Health grant R01-HL148348 (JES), National Institutes of Health (NIH) F31R03AA030408 (to J.L.S) a Washington University in St. Louis Rheumatic Diseases Research Resource-Based Center grant NIH P30AR073752 (to KJL), a National Institutes of Health grant R35-HL161185 (KJL), a Leducq Foundation Network grant #20CVD02 (to KJL), Burroughs Wellcome Fund grant 1014782 (to KJL), Children’s Discovery Institute of Washington University and St. Louis Children’s Hospital grant CH-II-2015-462, Ch-II-2017-628, PM-LI-2019-829 (to KJL), and a Foundation of Barnes-Jewish Hospital grant 8038-88 (to KJL)

## Competing Interests

SPC is an uncompensated ancillary Scientific Advisory Board member of Who We Play For, Inc. (WWPF), and a Scientific Advisory Board member for Rejuvenate Bio. SPC has a Sponsored Research Agreement (SRA) between WWPF and the FSU Office for the Vice President of Research. SPC was a recipient of sponsored research agreements from Pfizer, Rejuvenate Bio, and Implicit Bioscience. KJL is on the Advisory Board for Medtronic and is a recipient of sponsored research agreements from Amgen, Novartis, Implicit Bioscience and Kiniksa. JES is a consultant for Rejuvenate Bio, Implicit Bioscience and Rocket Pharmaceuticals. JES and Angeliki Asimaki hold–a US Patent (US Patent 10,317,417) for the use of buccal cells in the diagnosis of arrhythmogenic cardiomyopathy. KJL and JES are inventors in patent applications PCT/AU2023/051026 and PCT/AU2023/051027 both assigned to Implicit Bioscience Ltd.

## Supplemental Material

## Supplemental Methods

## Supplemental Figures S1-11

## Notes

### Competing Interest Statement

The authors have declared no competing interest.

## References

1. Scheel PJ, Murray B, Tichnell C, James CA, Tandri H, Calkins H, et al. Arrhythmogenic Right Ventricular Cardiomyopathy Presenting as Clinical Myocarditis in Women. Am J Cardiol 2021;145:128–134. doi:10.1016/j.amjcard.2020.12.090.

2. Corrado D, Basso C, Judge DP. Arrhythmogenic Cardiomyopathy. Circ Res 2017;121:784–802. doi:10.1161/CIRCRESAHA.117.309345.

3. Austin KM, Trembley MA, Chandler SF, Sanders SP, Saffitz JE, Abrams DJ, et al. Molecular mechanisms of arrhythmogenic cardiomyopathy. Nat Rev Cardiol 2019;16:519–537. doi:10.1038/s41569-019-0200-7.

4. Thiene G, Nava A, Corrado D, Rossi L, Pennelli N. Right Ventricular Cardiomyopathy and Sudden Death in Young People. New England Journal of Medicine 1988;318:129–133. doi:10.1056/NEJM198801213180301.

5. Al-Aidarous S, Protonotarios A, Elliott PM, Lambiase PD. Management of arrhythmogenic right ventricular cardiomyopathy. Heart 2024;110:156–162. doi:10.1136/heartjnl-2023-322612.

6. Cipriani A, Mattesi G, Bariani R, Cecere A, Martini N, De Michieli L, et al. Cardiac magnetic resonance imaging of arrhythmogenic cardiomyopathy: evolving diagnostic perspectives. Eur Radiol 2022;33:270–282. doi:10.1007/s00330-022-08958-2.

7. Wu I, Zeng A, Greer-Short A, Aycinena JA, Tefera AE, Shenwai R, et al. AAV9:PKP2 improves heart function and survival in a Pkp2-deficient mouse model of arrhythmogenic right ventricular cardiomyopathy. Communications Medicine 2024;4:38. doi:10.1038/s43856-024-00450-w.

8. Shiba M, Higo S, Kondo T, Li J, Liu L, Ikeda Y, et al. Phenotypic recapitulation and correction of desmoglein-2-deficient cardiomyopathy using human-induced pluripotent stem cell-derived cardiomyocytes. Hum Mol Genet 2021;30:1384–1397. doi:10.1093/hmg/ddab127.

9. Olson EN. Toward the correction of muscular dystrophy by gene editing. Proceedings of the National Academy of Sciences 2021;118. doi:10.1073/pnas.2004840117.

10. Campian ME, Verberne HJ, Hardziyenka M, de Groot EAA, van Moerkerken AF, van Eck-Smit BLF, et al. Assessment of inflammation in patients with arrhythmogenic right ventricular cardiomyopathy/dysplasia. Eur J Nucl Med Mol Imaging 2010;37:2079–2085. doi:10.1007/s00259-010-1525-y.

11. Engel M, Shiel EA, Chelko SP. Basic and translational mechanisms in inflammatory arrhythmogenic cardiomyopathy. Int J Cardiol 2024;397:131602. doi:10.1016/j.ijcard.2023.131602.

12. Hawthorne RN, Blazeski A, Lowenthal J, Kannan S, Teuben R, DiSilvestre D, et al. Altered Electrical, Biomolecular, and Immunologic Phenotypes in a Novel Patient-Derived Stem Cell Model of Desmoglein-2 Mutant ARVC. J Clin Med 2021;10:3061. doi:10.3390/jcm10143061.

13. Chelko SP, Penna VR, Engel M, Shiel EA, Centner AM, Farra W, et al. NFĸB signaling drives myocardial injury via CCR2+ macrophages in a preclinical model of arrhythmogenic cardiomyopathy. Journal of Clinical Investigation 2024;134. doi:10.1172/JCI172014.

14. Chelko SP, Asimaki A, Andersen P, Bedja D, Amat-Alarcon N, DeMazumder D, et al. Central role for GSK3β in the pathogenesis of arrhythmogenic cardiomyopathy. JCI Insight 2016;1. doi:10.1172/jci.insight.85923.

15. Chelko SP, Asimaki A, Lowenthal J, Bueno-Beti C, Bedja D, Scalco A, et al. Therapeutic Modulation of the Immune Response in Arrhythmogenic Cardiomyopathy. Circulation 2019;140:1491–1505. doi:10.1161/CIRCULATIONAHA.119.040676.

16. Penna VR, Amrute JM, Engel M, Shiel EA, Farra W, Cannon EN, et al. Interleukin-1β Drives Disease Progression in Arrhythmogenic Cardiomyopathy. JACC Basic Transl Sci 2026;11:101542. doi:10.1016/j.jacbts.2026.101542.

17. Tsukamoto H, Takeuchi S, Kubota K, Kobayashi Y, Kozakai S, Ukai I, et al. Lipopolysaccharide (LPS)-binding protein stimulates CD14-dependent Toll-like receptor 4 internalization and LPS-induced TBK1-IKKɛ-IRF3 axis activation. J Biol Chem 2018;293:10186–10201. doi:10.1074/jbc.M117.796631.

18. Na K, Oh B-C, Jung Y. Multifaceted role of CD14 in innate immunity and tissue homeostasis. Cytokine Growth Factor Rev 2023;74:100–107. doi:10.1016/j.cytogfr.2023.08.008.

19. Mikuls TR, Levan TD, Sayles H, Yu F, Caplan L, Cannon GW, et al. Soluble CD14 and CD14 Polymorphisms in Rheumatoid Arthritis. J Rheumatol 2011;38:2509–2516. doi:10.3899/jrheum.110378.

20. Thiebaut R, Charpentier C, Damond F, Taieb A, Antoine R, Capeau J, et al. Association of Soluble CD14 and Inflammatory Biomarkers With HIV-2 Disease Progression. Clinical Infectious Diseases 2012;55:1417–1425. doi:10.1093/cid/cis708.

21. Sharygin D, Koniaris LG, Wells C, Zimmers TA, Hamidi T. Role of CD14 in human disease. Immunology 2023;169:260–270. doi:10.1111/imm.13634.

22. Suzuki A, Fukuzawa K, Yamashita T, Yoshida A, Sasaki N, Emoto T, et al. Circulating intermediate CD14++CD16+monocytes are increased in patients with atrial fibrillation and reflect the functional remodelling of the left atrium. Europace 2016:euv422. doi:10.1093/europace/euv422.

23. Rogacev KS, Cremers B, Zawada AM, Seiler S, Binder N, Ege P, et al. CD14++CD16+ Monocytes Independently Predict Cardiovascular Events. J Am Coll Cardiol 2012;60:1512–1520. doi:10.1016/j.jacc.2012.07.019.

24. Xu J-J, Liu K-Q, Ying Z-M, Zhu X-W, Xu X-J, Zhao P-P, et al. Effect of CD14 polymorphisms on the risk of cardiovascular disease: evidence from a meta-analysis. Lipids Health Dis 2019;18:74. doi:10.1186/s12944-019-1018-3.

25. Stanislawski MA, Lange LA, Raffield LM, Zakai NA, Meyer M, Ferrier K, et al. Soluble CD14 Levels in the Jackson Heart Study: Associations With Cardiovascular Disease Risk and Genetic Variants. Arterioscler Thromb Vasc Biol 2021;41. doi:10.1161/ATVBAHA.121.316035.

26. Garcia-Alonso L, Holland CH, Ibrahim MM, Turei D, Saez-Rodriguez J. Benchmark and integration of resources for the estimation of human transcription factor activities. Genome Res 2019;29:1363–1375. doi:10.1101/gr.240663.118.

27. Schaller TH, Batich KA, Suryadevara CM, Desai R, Sampson JH. Chemokines as adjuvants for immunotherapy: implications for immune activation with CCL3. Expert Rev Clin Immunol 2017;13:1049–1060. doi:10.1080/1744666X.2017.1384313.

28. Yu S, Balasubramanian I, Laubitz D, Tong K, Bandyopadhyay S, Lin X, et al. Paneth Cell-Derived Lysozyme Defines the Composition of Mucolytic Microbiota and the Inflammatory Tone of the Intestine. Immunity 2020;53:398–416.e8. doi:10.1016/j.immuni.2020.07.010.

29. Ren F, Cai X, Yao Y, Fang G. JunB: a paradigm for Jun family in immune response and cancer. Front Cell Infect Microbiol 2023;13. doi:10.3389/fcimb.2023.1222265.

30. Baitsch D, Bock HH, Engel T, Telgmann R, Müller-Tidow C, Varga G, et al. Apolipoprotein E induces antiinflammatory phenotype in macrophages. Arterioscler Thromb Vasc Biol 2011;31:1160–1168. doi:10.1161/ATVBAHA.111.222745.

31. Akatsu C, Shinagawa K, Numoto N, Liu Z, Ucar AK, Aslam M, et al. CD72 negatively regulates B lymphocyte responses to the lupus-related endogenous toll-like receptor 7 ligand Sm/RNP. Journal of Experimental Medicine 2016;213:2691–2706. doi:10.1084/jem.20160560.

32. Liu H, Wang W, Shen W, Wang L, Zuo Y. ARHGAP24 ameliorates inflammatory response through inactivating Rac1/Akt/NF-κB pathway in acute pneumonia model of rat. Ann Transl Med 2020;8:1289–1289. doi:10.21037/atm-20-5000.

33. Dieterlen M-T, John K, Reichenspurner H, Mohr FW, Barten MJ. Dendritic Cells and Their Role in Cardiovascular Diseases: A View on Human Studies. J Immunol Res 2016;2016:1–13. doi:10.1155/2016/5946807.

34. Bibi S, Zeng W, Zheng P, Mousavi Mehmandousti SM, Jin T. Discoidin domain receptor 1 as a potent therapeutic target in solid tumors. HLife 2024;2:454–466. doi:10.1016/j.hlife.2024.01.003.

35. Friedman DJ, Kizerwetter M, Belmonte P, Rajcula M, Theodore K, Kim Lee HS, et al. Cutting Edge: Enhanced Antitumor Immunity in ST8Sia6 Knockout Mice. The Journal of Immunology 2022;208:1845–1850. doi:10.4049/jimmunol.2101165.

36. Zhang H, Meng F, Chu C-L, Takai T, Lowell CA. The Src Family Kinases Hck and Fgr Negatively Regulate Neutrophil and Dendritic Cell Chemokine Signaling via PIR-B. Immunity 2005;22:235–246. doi:10.1016/j.immuni.2005.01.004.

37. Huang Q, Wang F, Hao D, Li X, Li X, Lei T, et al. Deciphering tumor-infiltrating dendritic cells in the single-cell era. Exp Hematol Oncol 2023;12:97. doi:10.1186/s40164-023-00459-2.

38. Ong S, Rose NR, Čiháková D. Natural killer cells in inflammatory heart disease. Clin Immunol 2017;175:26–33. doi:10.1016/j.clim.2016.11.010.

39. Soussi S, Maione AS, Lefèvre L, Pizzinat N, Iacovoni J, Gonzalez-Fuentes I, et al. Analysis of effector/memory regulatory T cells from arrhythmogenic cardiomyopathy patients identified IL-32 as a novel player in ACM pathogenesis. Cell Death Dis 2025;16:87. doi:10.1038/s41419-025-07364-y.

40. Maione AS, Pilato CA, Casella M, Gasperetti A, Stadiotti I, Pompilio G, et al. Fibrosis in Arrhythmogenic Cardiomyopathy: The Phantom Thread in the Fibro-Adipose Tissue. Front Physiol 2020;11. doi:10.3389/fphys.2020.00279.

41. Amrute JM, Luo X, Penna V, Yang S, Yamawaki T, Hayat S, et al. Targeting immune– fibroblast cell communication in heart failure. Nature 2024;635:423–433. doi:10.1038/s41586-024-08008-5.

42. Xu Y, Liu Y, Yang C, Kang L, Wang M, Hu J, et al. Macrophages transfer antigens to dendritic cells by releasing exosomes containing dead-cell-associated antigens partially through a ceramide-dependent pathway to enhance CD4(+) T-cell responses. Immunology 2016;149:157–171. doi:10.1111/imm.12630.

43. Zhou J, Zhang S, Guo C. Crosstalk between macrophages and natural killer cells in the tumor microenvironment. Int Immunopharmacol 2021;101:108374. doi:10.1016/j.intimp.2021.108374.

44. Yang B, Qiao Y, Yan D, Meng Q. Targeting Interactions between Fibroblasts and Macrophages to Treat Cardiac Fibrosis. Cells 2024;13. doi:10.3390/cells13090764.

45. Kadyrov FF, Koenig AL, Amrute JM, Dun H, Li W, Weinheimer CJ, et al. Hypoxia sensing in resident cardiac macrophages regulates monocyte fate specification following ischemic heart injury. Nature Cardiovascular Research 2024;3:1337–1355. doi:10.1038/s44161-024-00553-6.

